# Neurons from pre-motor areas to the Mushroom bodies can orchestrate latent visual learning in navigating insects

**DOI:** 10.1101/2023.03.09.531867

**Authors:** Antoine Wystrach

**Affiliations:** Centre de Recherches sur la Cognition Animale, CBI, CNRS, Université Paul Sabatier, Toulouse, France

**Keywords:** insects, navigation, vision, latent learning, homing, memory, mushroom bodies, central complex, lateral accessory lobes, neural modelling

## Abstract

Spatial learning is peculiar. It can occur continuously and stimuli of the world need to be encoded according to some spatial organisation. Recent evidence showed that insects categorise visual memories as whether their gaze is facing left vs. right from their goal, but how such categorisation is achieved during learning remains unknown. Here we analysed the movements of ants exploring the world around their nest, and used a biologically constrained neural model to show that such parallel, lateralized visual memories can be acquired straightforwardly and continuously as the agent explore the world. During learning, ‘left’ and ‘right’ visual memories can be formed in different neural comportments (of the mushroom bodies lobes) through existing lateralised dopaminergic neural feedback from pre-motor areas (the lateral accessory lobes) receiving output from path integration (in the central complex). As a result, path integration organises visual learning ‘internally’, without the need to be expressed through behaviour; and therefore, views can be learnt continuously (without suffering memory overload) while the insect is free to explore the world randomly or using any other navigational mechanism. After learning, this circuit produces robust homing performance in a 3D reconstructed natural habitat despite a noisy visual recognition performance. Overall this illustrates how continuous bidirectional relationships between pre-motor areas and visual memory centres can orchestrate latent spatial learning and produce efficient navigation behaviour.

## MAIN

Insect navigators such as ants, bees and wasps rapidly learn the visual surroundings to navigate efficiently to places of interest such as the nest or food sources ^1^. These long-term visual memories are formed in a brain area called the **Mushroom Bodies (MBs)**^2,3^. The neural circuitry of the MBs is ideally suited to encode, store and compare arbitrary input (visual, olfactory, or other) in a way that enables a visually navigating insect to learn and then assess whether the visual scene currently perceived is familiar or not ^4,5^. After learning, the MBs output ‘familiarity signals’ that can be used for guidance along familiar routes or back to place of interest ^4,6,7^. However, how learning is orchestrated in the first place is unclear.

As during experimental conditioning, experiencing an event bearing an innate positive or negative valence (so called the US in learning theory) can trigger the learning of the surrounding visual scenery (which can be viewed as the CS). For instance, experiencing sucrose at a feeder location will trigger visual learning events that are useful to return to this rewarding place ^8–10^. Inversely, ants experiencing a negative event such as falling into a pit-trap will memorise the visual scenes experienced just before falling as aversive; and hence avoid this region of the world in the subsequent trips ^11,12^. These examples involve a reinforcer (reward or punishment) and thus fall under the umbrella of ‘reinforcement learning’. However, spatial learning also occurs in the absence of distinctive reward or punishment, for instance, when exploring the world. Indeed, navigating insects tend to learn continuously: weather along routes, around their nest (during so-called learning walks or learning flight) but also when at novel albeit quite neutral locations ^13–15^. This tendency to learn continuously when exploring the world is shared with other navigating animals too, and has been dubbed ‘latent learning’ in opposition to ‘reinforcement learning’ in learning theory ^16^

Whether ‘latent’ or ‘reinforcement-based’, spatial learning implies that the stimuli of the world are encoded according to some spatial organisation. For navigating ants, it has been suggested that learning of the visual surroundings may happen only when the ant is facing specific directions of interests, such as when its gaze is oriented towards the goal ^17–21^ the anti-goal ^6,22,23^ or along their route direction ^4,7,24–26^. At the naïve stage, this directional information can be provided by **path integration (PI)**. PI continuously provides the insect’s current position relative to its goal, whether the nest or a food source ^27^. It has thus been suggested that path integration is used to both enable the physical alignment of the insect’s body and gaze towards its goal (or anti-goal) and trigger a visual learning event at such appropriate times ^19,28^. PI is computed in a brain area called the **Central complex (CX)**, the seat of the insect representation of directions ^29–32^. However, how the right information from the path integrator is mediated to the MBs for orchestrating visual learning remains entirely unknown^33^. Recently, it was shown that ants (and likely wasps ^34^) may not learn views specifically when facing the goal or anti-goal direction, but when facing left and right from their goal ^35^. Left-to-the-goal and right-to-the-goal memories explains why ants can recognise egocentric views when misaligned with their goal and trigger the appropriate turning commands ^34–36^, but summon an explanation for how left/right categorisation of visual memories is achieved neurally during learning.

Here we show how such categorisation can be achieved neurally and continuously, providing a mechanistic explanation for spatial, latent learning. We analysed the movements of ants displaying learning walks around their nest, and used computational modelling to show that existing neural feedback from pre-motor area **(the lateral accessory lobes, LAL**) receiving the path integration output from the CX, can organise the formation of these lateralized visual memories in the MBs. Our biologically constrained architecture shows that learning events can then be achieved randomly or continuously; literally sparing both the need to ‘control the timing of learning’ as well as the need to align the agent’s body in any particular direction. Remarkably, the MBs can support continuous learning of thousands of views without suffering memory overload, because only novel information recruits new synapses. After learning, the architecture can produce remarkably robust homing performance in reconstruction of complex natural habitats ^37^, as observed in homing ants ^38–41^.

### Ants look in all directions during learning walks

During learning walks, naïve ants display meandering trajectories around their nest, often exploring different directions multiple times before venturing further ^21^. Artificially restraining these exploratory movements (in both time and space) reduces the ants’ subsequent navigational performance based on terrestrial cues, showing that they do learn the scenery during these exploratory behaviours ^39,41^. At a finer scale, these meandering trajectories are interspaced with regular slowing down up to complete halts, producing behaviours so-called votes or scans, whose expression varies across species and individuals^42,43^. Slowing down and pausing helps the ants obtain a stable (and thus non-blurry) view and therefore surely contributes to visual learning. This is corroborated by the fact that these pauses are typically displayed in situation when learning is needed ^9,11,15,19,44–48^. It was shown in some species that ants tend to display longer pauses when their gaze is aligned towards the nest ^19,43^ or anti-nest^22^ directions.

Here, by analysing the learning walks recorded at high frame rate of two species of ants (*Myrmecia croslandi* and *Melophorus bagoti*), we found no systematic association between such pauses and some particular nest-centred or allocentric directions (Fig. 1c, Extended data Fig. 3). In contrasts, ants (pausing or not) exposed their gaze to a large, homogenous diversity of directions and locations around their nest (Fig. 1c, Extended data 3). While we acknowledge that ants may sometimes pause for a longer period of time while looking in the direction of their goal, what is clear is that learning walks are rather optimised to collect a diverse sample of views in all directions. This is in line with previous works showing that pauses and scans are not tightly controlled by the relation with the ant and its environment, but rather are the result of ‘blind’ internal motor processes such as the continuous production of regular oscillations in the ant’s angular and forward speed ^22,49^ as well as the random triggering of pauses ^42^. This stochasticity is further highlighted by the great variability in the expression of learning walks observed across individuals ^50^.

**Figure 1.**
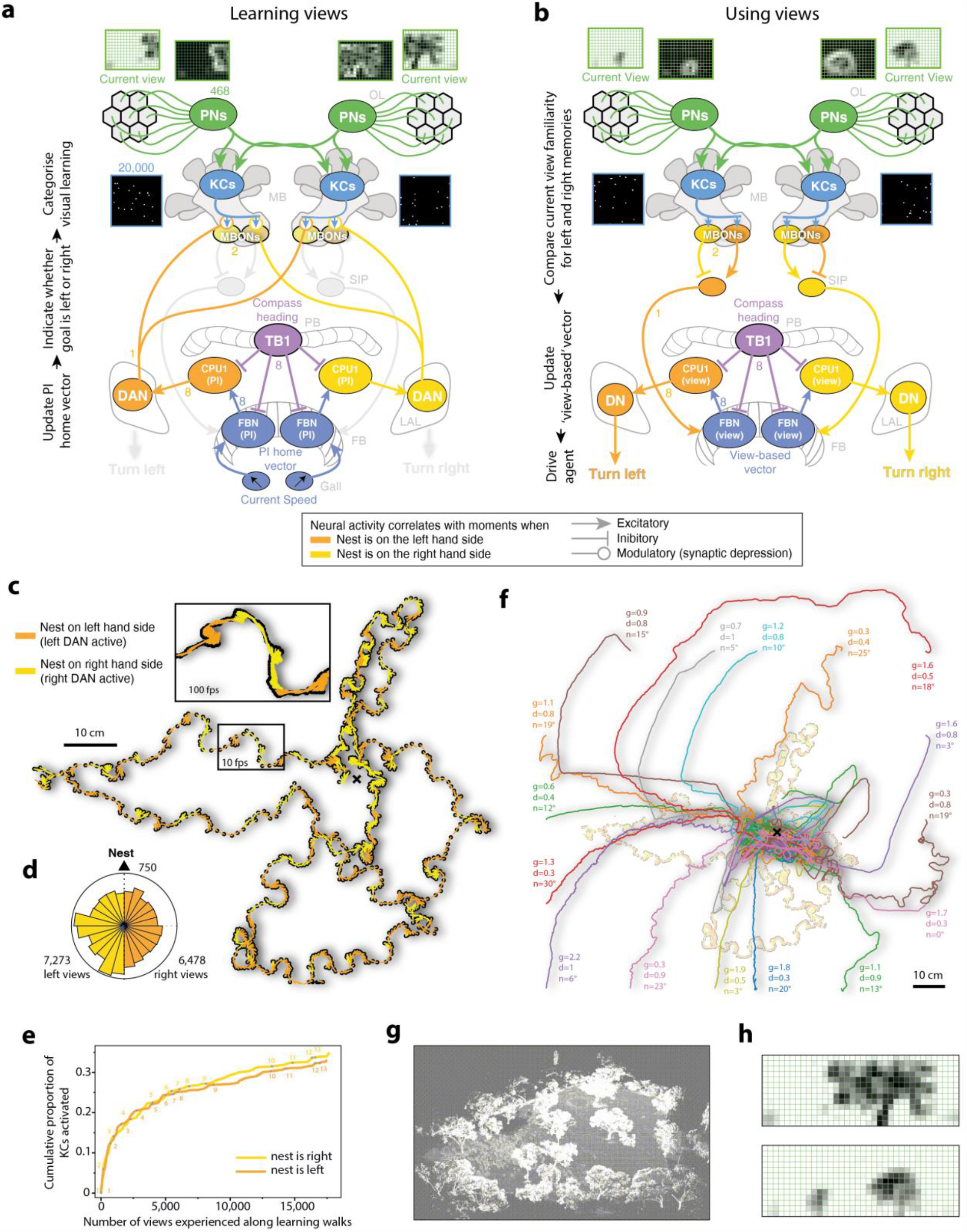
A Mushroom Bodies and Central Complex circuit produces robust visual navigation. **a,b**. Schematic of the model’s functional circuitry. The agent’s current panoramic view results from its position in the reconstructed world (**g**), down-sampled at 10 °/pixel (**h**). Projection Neurons (PNs) sample the whole visual field and form random connections with Kenyon Cells (KCs), resulting in a pattern of KC activity highly specific to the current view. Numbers on the left indicate numbers of neurons. See Extended data 1 for details. **a**. During learning, the central complex is updating a path integration (PI) home vector by integrating current speed and compass heading information as in ^32^. The output of the CX in the left (or right) hemisphere’s LAL, which correlate with the time when the nest is on the left- (or right-) hand side – and thus can be used to drive left (or right) turns to home by PI – are used instead to drive dopaminergic neurons (DAN) projecting to the MBs to categorise visual learning. DANs’ activity triggers memory formation by synaptic depression of the currently active KCs outputs on the associated MBS output neurons (MBONs). **b**. After learning, familiar views differentially activate MBONs according to the KC-MBON synaptic strengths established during learning. MBONs’ signals are integrated in the SIP (Superior Intermediate Protocerebrum) as an opponent-like process (Le Möel and Wystrach, 2020) providing a measure of the likelihood of having the nest on the left or on the right, that is independent of the overall level of visual familiarity ^35^. These lateralized signals then project to the CX (as shown in figure 3), literally updating a ‘view-based vector’ representation. Motor control is effected by the usual CX circuitry based on the current compass heading (as for PI), resulting in the agent performing turns. (OL: Optic-Lobes, MB: Mushroom-Bodies, SIP: Superior-Intermediate-Protocerebrum, PB: Protocerebral Bridge, FB: Fan-shaped-Body, LAL: Lateral Accessory Lobe). **c**. Example of two consecutive learning walks displayed by an individual *Myrmecia crosslandi* ant (data courtesy of Jochen Zeil) and used by the agent for learning views. In this example, the agent sampled the world at 100 fps (see inset for realistic representation of sampled views’ positions) approximating the ants visual flicker fusion frequency and thus assuming continuous learning (Extended data Fig. 2 shows other training conditions). **d**. Circular histogram of the number of views experienced along the learning walks (**c**) according to their orientation relative to the nest, showing that the ant exposed its gaze in all directions relative to the nest (750 indicate the scale at the circle rim). **e**. Cumulative proportion of KCs activated at least once along 13 consecutive learning walks (Extended data Fig. 2). The tendency to plateau explains how continuous learning can be supported without memory saturation. **f**. Paths realised by agents using views (**b**) in closed-loop with the environment to home from novel release locations around the nest, after learning (**a**) along two learning walks (**c**). The agents display efficient homing and nest search across a range of randomly chosen parameter values (g: motor gain; d: FBN decay; n: motor noise, see parameter description and Extended data Fig. 1 for detailed explanation). **g**. Visual reconstruction of the Myrmecia ants’ natural environment ^37^ used in the current simulation (represented as a points cloud for clarity). **h**. Example of views drawn from the reconstructed world, down-sampled at 10 °/pixel to ensure that ant resolution is not overestimated.

### Visual learning in the MB lobes can be organised by lateralized dopaminergic feedback from pre-motor areas

Previous works have suggested that ants and wasps categorise views as whether their gaze is oriented towards the left vs. right in relation to the nest heading direction ^34,35^; but if ants look in all directions during their learning walks, how do they achieve such a left vs. right categorisation?

We realised that due to the Path integration – the ability to integrate compass and distance information to keep track of the nest relative position ^51^ – the output of the **Central complex (CX)** to the **Lateral Accessory Lobe (LAL)** provides the desirable information: the left (or right) LAL’s hemisphere activity correlates with moments when the nest relative position is on the left (or right) of the ant current heading direction ^32,52^. The LAL are pre-motor areas sending steering commands to neurons descending to the thorax ^49,53^. When PI is controlling guidance for homing, these lateralized output signals are used to trigger ‘turn left’ and ‘turn right’ compensatory motor commands to align the insect’s body towards its nest, hence resulting in homing behaviour. However, when PI is not used to home –such as when ant display a learning walk – we reason that the LAL’s output could nonetheless be used, not for steering, but to control ‘internally’ whether the current view should be categorised as left or right from the goal during learning (Fig. 1a).

Interestingly, the insect brain possesses the perfect neural candidate to do so: direct dopaminergic projections from the LALs to the MBs lobes ^54^(Fig. 1a), that is, where long-term visual memories are formed due to dopamine release ^55,56^. The left and right dopaminergic feedback from the LAL –which could thus indicate in real time whether the nest is left or right of the current heading – project to different compartments of the MBs lobes (see Fig. 4D of ^54^); so that ‘left-to-the-goal’ and ‘right-to-the-goal’ memories could be formed in the input synapses of different **MB output neurons (MBONs)**, literally updating separated memory banks as the individual explores the scene: the ‘left-to-the-goal’ memory bank is updated when the nest direction is on the left side of the insects current facing direction, and vice versa. Note that other neural candidates could equally achieve the desired LAL-to-MBs learning signals, albeit indirectly. For instance, some feedback from pre-motor areas modulate dopaminergic neurons that in turn, trigger synaptic modulation in the MBs lobes ^57,58^.

### Homing can be achieved through opponent lateralized signals from the MBs to the CX

Once the views are memorised and categorised as left vs right, subsequent homing – based on these learnt views – requires to convey the familiarity signals from the MB lobes to the LAL for steering. We have strong behavioural and neurobiological evidence to constrain our explanation of how this might happen.

1. Connectomic ^52^ and experimentation in ants ^35^ shows that navigation based on learnt views is achieved indirectly, by updating a goal heading compass direction, likely in the fan-shaped body of the CX, which in turns control steering in the LAL. Interestingly, this seems to work only if the familiarity signals sent to the CX are decorrelated between the left and right hemispheres, with the signal from the left (and right) hemisphere indicating when the animal current heading is biased towards the left (or right) compared to the goal direction ^35^.
2. Connectomic suggests that the MBs lobes send the familiarity signals to the CX either directly ^59^, or through one relay in the dorsal brain areas (such as the SIP) ^52^, where most MBONs converge ipsilaterally (^59^. Such a relay notably enables the integration of antagonist MBON (and other) signals conveying opposite valences through simple inhibition ([good - bad], or [bad – good]) ^60^. This produces an opponent-like process which improves the estimation of the valence of the current situation and appears to be at play during visual navigation in ants ^6,23^.
3. Some dopaminergic neurons from the left and right LAL – which are thought here to organise learning at the first place (see previous section) – both project bilaterally in the MB lobes in a symmetrical manner (see Fig.4D ^54^, suggesting that both left and right hemispheric MBs encode both left and right view memories in two different MBs lobes compartments each (4 different compartments in total) (Fig. 1a ‘DAN’).

We realised here that these three points naturally converged into one picture (Fig.1a,b), which produces a set of predictions:

1. In each hemisphere, ‘left and right memories’ are formed in MBs compartment conveying opposite valence, in a symmetrical manner (Fig. 1a ‘DAN’, point 3 above).
2. During homing, the resulting ‘left and right familiarity signals’ (mediated by different MBONs) are then integrated ipsilaterally (in the dorsal brain area relay) as an opponent process (Fig. 1b ‘MBON’, point 2 above).
3. Due to the symmetry, this integration is achieved in an opposite manner in each hemisphere (Left – Right familiarity in the left hemisphere; and Right – Left familiarity in the right hemisphere) Fig. 1b ‘SIP’).
4. Both opponent signals are then sent to the CX ipsilaterally, providing the desired uncorrelated left and right familiarity input to the CX (point 1 above) (Fig. 1b, ‘FB’).

### A robust circuit for navigation in noisy visual environments

To proof-test the validity of this circuit as a whole, we modelled both the MBs and the CX based on connectomic data as achieved before (for MB: ^4^, for CX: ^32^), and coupled these two brain regions using the mentioned LAL-to-MBs connections for learning views, and MB-SIP-CX connections for using views (Fig. 1, Extended data Fig. 1). This circuit was implemented into an agent with ant-eye-like resolution (10°/pixel, Fig.1.h), immersed in a realistic VR reconstruction of Myrmecia ants’ natural habitat ^37^ (Fig.1g).

For learning, we let the agent reproduce recorded natural learning walks of a Myrmecia ant (Fig. 1c, Extended data Fig.2; data provided by Jochen Zeil), with the outputs of its path integrator (computed in the CX) to the left and right LALs (Fig.1c, yellow and orange) driving the dopaminergic feedbacks that control learning. Learning in the MB is achieved following what is observed in insects: by depressing the currently active KCs’ output synapses onto the MBONs, only in the MBs compartments where the associated dopaminergic feedback is concomitantly active ^55,61^. After completion of the learning walk, we transferred the agent deprived of PI (i.e. as a so called Zero-Vector agent) to a new location in the world and let the visual familiarity based on the left and right memories – as outputted by the MBONs – drive the two FB inputs to update the desired heading direction in the CX. The CX, in turn, outputs left and right turning commands to the LAL, which drives the agent (Fig.1,b, Extended data Fig.1). This produces a remarkably robust homing ability, as well as a tight search around the nest without the need to fine tune parameters (Fig. 1f, Extended data Fig. 2), providing further credibility to this circuit.

### When to learn? No need to bother

Interestingly, our model shows that the timing and positions at which the view is sampled along learning walks are not important. The model can afford to learn sporadically, at random position or continuously (it is operational whether it uses 90 views or 30,000 views for learning) (see Extended data Fig. 2), bypassing the need for additional mechanisms that controls ‘when to learn’ or ‘how to align the body to learn’. Memory load is not a problem either: even when learning continuously (i.e., here at 100fps), 20,000 KCs proved largely sufficient to store the information from the large amount of views (e.g., 13638 views) experienced along a learning walks sampled at high frame rate (100 fps). Saturation is prevented because additional memory space is used only when significantly novel views are perceived: already learnt views activate KCs that have already switched their synaptic output, and thus yield no further change. Acquiring visual memories in this way results in a steep learning curve at first, but then spontaneously plateaus as the explored regions around the nest become familiar (Fig. 1e). In addition, our model vastly underestimates the ants’ (and hymenopteran navigators in general) memory capacity (>100,000 KCs ^62,63^ and >100 MBONs ^64^), ignores pruning principles that can increase effective memory capacity, bypasses any visual pre-processing that can compress information, and reduces the complexity of synaptic connections to simple binary connections between neurons. Hence, we are confident that learning views around the nest in such a way should occupy only a fraction of the real ants’ memory capacity.

### General discussion

We have shown how existing neural connections between the CX (computing path integration), the LAL (a pre-motor area) and the MB (seat of the visual memories), could enable an insect to orchestrate spatial learning. An agent equipped with this circuit can quickly learn the visual surrounding and subsequently home from novel locations using very low-resolution views (10°/pixel, Fig. 1h) of its natural environment (Fig. 1). Several differences contrast this circuit from previous accounts.

First, views are encoded egocentrically (i.e. recognition is view-point dependant), which is in adequation with most of the literature ^24,65–69^ and contrast with recent models who assumed that visual memories can be rotationally invariant (recognition is independent of the gaze orientation) ^7,70^. The latter requires computational steps, such as Fourier transforms, which potential implementation in insect circuits remains unknown. Instead, assuming an egocentric encoding enables to remain faithful to the known insect neural circuits and corroborates the behavioural evidence that ants’ visual scene recognition requires the insect to align its gaze as during training ^66,71–73^ as well as the regular need of insects to scan multiple directions ^21,42,49,74,75^.

Second, our circuit is built on the evidence that memorised views are categorised as oriented left vs. right rather than towards or away from the goal ^34,35^. This was key, as this left/right information can be directly provided by the LAL during learning, due to the mechanism of path integration in the CX.

Third, it was previously assumed that path integration serves as a scaffold for visual learning through behaviour: the insect would use PI to physically align its body in a direction of interest (towards or away from the nest, or along a route) and then memorise the view perceived at this precise moment ^19,20,43^. Here, PI still serves as a scaffold for visual learning, but does so ‘internally’ without the need to express through behaviour. As a consequence, views can be learnt continuously while the insect freely explores the world using other mechanisms. With this in mind, ants learning walks ^21^ and early meandering trajectories ^50,76,77^ appear optimised to sample views quickly in as many locations and orientations around the nest as possible (Fig1 c,d; Extended data Fig. 3). Also, since all views becomes useful whatever their orientation, it explains how views acquired during outbound trips can be equally categorised effectively to serve subsequent homing ^46,73,78^, as well as the ants’ ability to recognise views whatever their body orientation when on highly familiar regions ^35^.

Fourth, navigation robustness arises here from multiple reasons. As shown before, visual recognition in the MBs is intrinsically noisy, partly because of the way memories are compressed in the KCs, and partly because of the proximal clutter (blade of grasses, etc…) that an insect may encounter when walking on the floor or flying through bushes ^4,5^. Part of this noise is alleviated by having the noisy MBs familiarity signal being send to the CX (as in^7,25,33,35^, which acts as a directional buffer due to its much more stable heading encoding based on multiple sources of compass ^30^. The current circuit provides an additional level of robustness, through redundancy, as each view is simultaneously compared to four memory banks in parallel (Fig.1b, ‘MBONs’). Although the input from each eye is sent to both left and right hemispheric MBs ^63^, the random pattern of connectivity in the KCs input ^79–81^ makes each MBs different. As a result, this architecture literally provides four independent assessments of whether the insect’s current heading is too much on the left vs. the right of its goal direction. Additional assessments, for instance based on recent vs. longer-term memories, may likely recruit more MBs compartments and make the recognition process more redundant.

Finally, the dopaminergic internal feedback from the LAL – necessary to categorise learning into different MB lobes compartments – also predicts a strong link between locomotion and activity in the MB, with rapid shifts of dopaminergic activity across the MB lobes as the insects is moving in the world. This corroborates recent observations in drosophila that dopamine release in the brain strongly correlates with the animals’ movements ^58,82^, and indeed, shifts dynamically across the MBs lobes ^57,83^. Overall, this supports the view that the MBs provide an active coding that operates in a tight closed-loop with the ongoing behaviour.

Which actual neurons convey the desired PI information from the LAL to the MBs to categorise visual learning in ants, wasps or bees remains to be identified, but given the existence of such direct (or indirect) connections in various insects that are not central place foragers ^54^, we can easily envision how the ability to perform path integration may have enabled the evolution of the ability to home using learnt views.

## MATERIALS AND METHODS

### Neural model

We used a simple agent-based model in a closed loop in a 3D virtual environment. All simulations were performed in Python 3.x. ^6^. The neural model connectivity is described in Extended data Fig. 1, and code is available on request.

### Parameters description

#### Motor noise

at each time step, a directional ‘noise angle’ is drawn randomly from a Gaussian distribution of ± SD = motor noise, and added to the agent’s current direction.

#### Memory decay

proportion of Fan-shaped Body Neurons (FBN, see extended Fig.1 for details) activity lost at each time step: For each FBN: Activity_(t+1)_ = Activity_(t)_ x (1 - memory decay). This corresponds to the speed at which the memory of the vector representation in the FBN decays. A memory decay = 1 means that the vector representation in the FBN is used only for the current time step and entirely overridden by the next inputs. A memory decay = 0 means that the vector representation acts as a perfect accumulator across the whole paths (as for Path Integration), which is probably unrealistic.

#### Motor gain

Sets the gain to convert the motor neuron signals (see extended Fig.1 for details) into an actual turn amplitude (turn amplitude = turning neuron signal × gain). Note that here, the motor gain is presented across orders of magnitude. One order of magnitude higher means that the agent will be one order of magnitude more sensitive to the turning signal.

### 3D world and view rendering

The virtual environment used in our model was generated by the software Habitat3D ^84^, an open-source tool which provides photorealistic meshes of natural scenes from point clouds acquired with help of a LiDAR scanner (IMAGER 5006i). This environment is mapped on the habitat of *Myrmecia* ants from Canberra, Australia ^37^. The mesh spans a diameter of around 65 metres and features large eucalyptus trees and the distant panorama cues (Fig. 1g). This dataset can be found on the Insect Vision webpage (https://insectvision.dlr.de/3d-reconstruction-tools/habitat3d). For speed optimization purposes, we down-sampled the originally high-resolution mesh with the open-source software Blender into a lower number of vertices; the rendering was then realized in OpenGL, with the Python libraries Plyfile and PyOpenGL. This 3D model enabled us to render panoramic views from any location as a 360-degree picture. We chose input only the blue channel of the RGB scene, resulting in only one luminance value per pixel. Also, the skybox was a pure blue uniform colour. That way, as with UV in natural scenes ^85,86^, blue provides a strong contrast between the sky and the terrestrial scene. This approximates the type of visual information used by navigating ants ^87,88^. Views were cropped vertically so that the bottom, floor-facing part was discarded. Finally, views were downsampled at 10°/pixel (see Fig. 1h), and we extracted the edges by subtracting for each pixel the summed value of its 8 neighbours, mimicking lateral inhibition across ommatidia. As a result, the visual information that the model receives is a small rectangular matrix of single-channel, floating point values representing the above-horizon panorama (see Fig. 1h).

### Learning walks trajectories analysis

We analysed learning walks trajectories of *Myrmecia croslandi* ants recorded at 100fps (courtesy of Jochen Zeil, from a data set used in ^22^) and *Melophorus bagoti* ants recorded at 300fps (from a data set used in ^15^). Trajectories were analysed using Matlab (R2016b Matworks). Code and data are available on request.

## List of abbreviations

PI: Path integration
MB: Mushroom Body
CX: Central complex
LAL: Lateral accessory lobes
MBON: Mushroom body output neuron
DAN: Dopaminergic neuron

## ACKNOLEDGMENTS

I am grateful to Florent Le Moël for setting up the python environment to run agent-based-simulations, as well as to Rüdiger Wehner, Paul Graham, Tom Collett and Michael Mangan for their useful feedbacks on an earlier draft of the manuscript. This study was funded by the European Research Council 759817-EMERG-ANT ERC-2017-STG

---

**Extended data figure 1.**
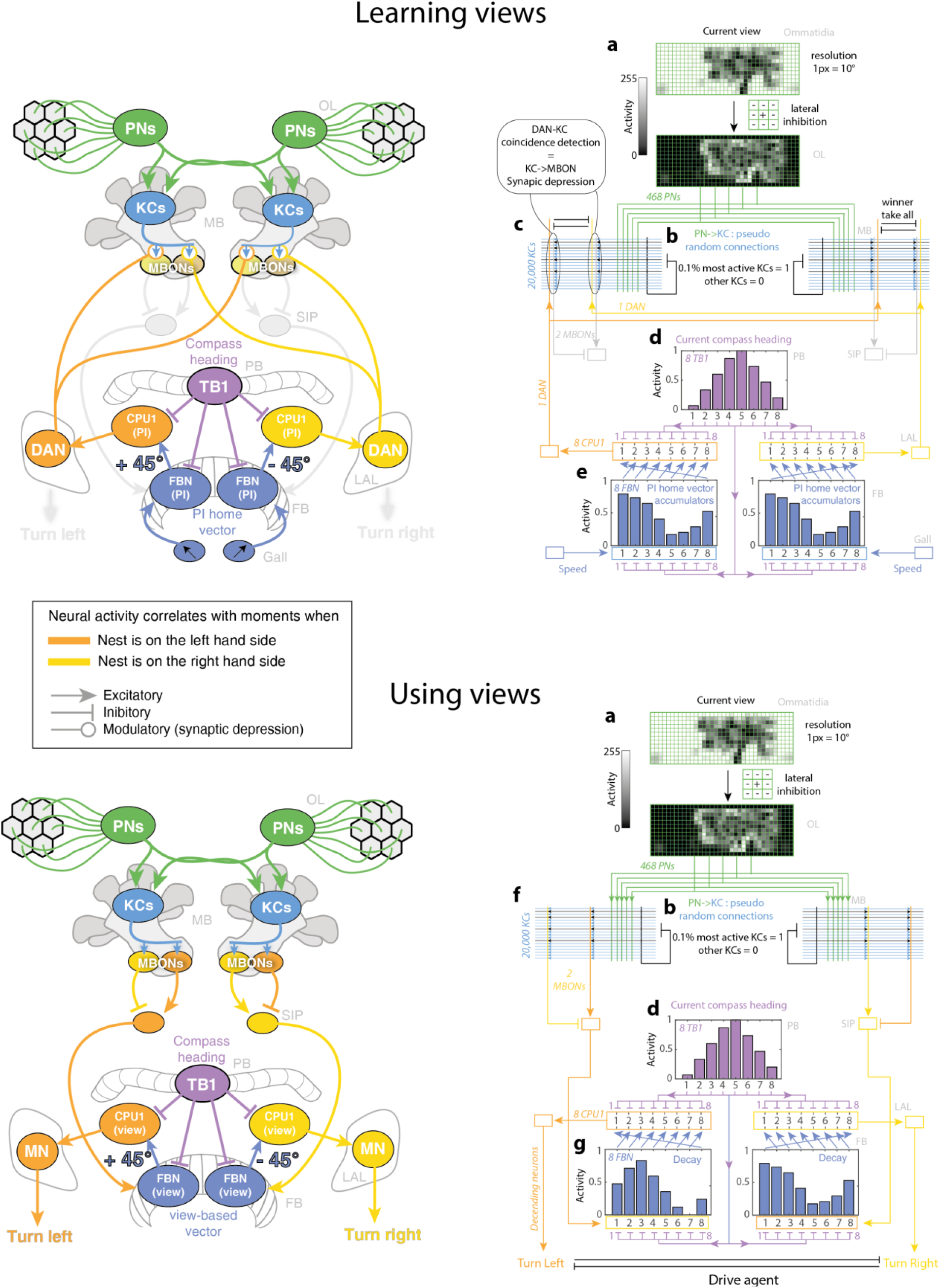
Details of the MB-CX model’s circuitry. Circuitry used for learning views (top), and using views to drive the trained agent (bottom). Left panels show the scheme as presented in figure 5, and right panels show the corresponding detailed circuitry. **a**. The agent’s current view (360° panoramic, with 90° above and 40° below horizon) is extracted from the reconstructed world at 10°/pixel, so 36 × 13 pixels = 468 cells. Activity of the cells correspond to the pixel light intensity (from 0 to 255) and could be seen as representing the cells’ firing rate. The view is processed through lateral inhibition between neighbouring cells: cell activity = cell activity – (∑ (all neighbouring cells activity) / number of neighbouring cells). This well-known early visual pre-processing makes cells respond to contrasted edges in the view, which is necessary for the downstream Kenyon Cells (KCs) to encode view specificity. **b**. Each view cell projects (via Projection Neuron, PN) to both hemispheres’ Mushroom Bodies (MB), where it makes pseudo-random connections with KCs: we set each KC to connect to 4 randomly chosen PNs, roughly matching what is observed in insects. We chose 20,000 KCs per hemisphere, which underestimates the number of KCs in ants (> 100,000). At each time step, the 0.1% KCs with the strongest input (i.e., the sum of the 4 PNs activities connecting to the KC, which can be seen as the KC’s dendritic excitatory postsynaptic potential) activity would be set to 1 (reflecting one action potential), the other KCs would be set to 0. This represents the effect of the inhibitory activity of APL-like-neurons (black neuron) across all KCs, ensuring that only a few KCs (the ones with strongest input activity) can fire an action potential at a time, as observed ^81,89,90^. **c**. We modelled (in each hemisphere) two compartments of the MBS lobes (surrounded by black ovals): both compartments are composed of 1 dopaminergic neuron (DAN) associated to 1 MBS output neuron (MBONs), mediating opposite valences as observed across insects ^91,92^. These antagonistic DANs engage in a winner-take-all competition (symbolised by the black reciprocal inhibition) so that only one kind is active at a time in each hemisphere, as observed in insects ^57^. Initially, all KCs connect to both MBONs with a synaptic weight of 1. At each time step, synaptic depression happens for the active DAN’s compartment mimicking coincidence detection ^55^: the KC-to-MBON weights of each currently active KCs is set to 0, and will stay so permanently (we did not wish to model forgetting). Due to the activity of the CX (see (e)), the DANs activity correlates with moments when the nest is left (orange DAN) or right (yellow DAN) relative to the current body orientation. **d**. Current compass direction is modelled in the protocerebral bridge (PB) as a bump of activity across 8 neurons forming a ring-attractor, as observed in insects ^52^. Each neuron responds maximally for a preferred compass direction, 45° apart from the neighbouring neurons (neuron 1 and 8 are neighbours). Change in the agent’s current compass orientation results in a shift of the bump of activity across the 8 neurons (we did not model how this is achieved from sensory compass cues (see ^30,93–95^ for studies dedicated to this matter). **e**. During learning, two representations of the Path Integration (PI) home vector are updated in the Fan-shaped Body Neurons (FBN) by integrating current speed and compass heading information (as in ^29,70^). Speed input activates all 8 FBN neurons equally, but simultaneous inhibition from the PB (see d) results in a negative imprinting of the current bump of activity (inhibition is effected between each paired neuron: 1 inhibits 1; 2 inhibits 2; etc…). FBN activity is sustained (given a slow decay), and thus acts as a PI home vector accumulator (Stone). Neurons, called CPU1 in some insects, compare each version of the home vector neurally shifted by 1 neuron (as if rotating the ring attractor representation by 45° clockwise or counter-clockwise depending on the hemisphere) with the current compass heading, resulting in an overall activity in the CPU1 (sum of the 8 CPU1) indicating whether the nest is rather on the left- (higher activity in the left hemisphere) or right-hand side (higher activity in the right hemisphere). This left/right differential activity – instead of driving the agent home – is integrated in a DAN connecting the LAL to the MBs (described in Fig. 4D of ^54^) and thus used to categorise visual learning (see c). **f**. The current view results in a specific pattern of KC activity (a), which activates MBONs differentially according to the weight of the KC-MBONs connections set during learning (c). For instance, views similar to the one experienced when the nest was on the left (orange DAN in (c), trigger KCs with KC-MBON weight set to 0 in this compartment, and thus will activate mostly the MBON of the other compartment. This differential activity between MBONs is integrated in the SIP (Superior Intermediate Protocerebrum) in each hemisphere, resulting in an opponent-like process providing a measure of the likelihood of having the nest on the left or right that is independent of the overall level of visual familiarity (similarly to ^6,23^). **g**. These lateralized signals from the SIP excite a dedicated set of FBN, literally updating a ‘view-based vector’ representation. The sustainability of such a ‘view-based vector’ depends on the FBN activity’s decaying rate, which can be varied in our model and has little incidence on the agent success (Extended Data Fig. 2, parameter decay). Motor control is effected using the same circuitry than for PI ^29,32^: the CPU1 neurons control descending motor neurons (MN), which difference in activity across hemispheres triggers a left or right turn of various amplitude, given a ‘motor gain’ that can be varied to make the agent more or less reactive (see parameter description). Numbers on the left indicate neuron numbers. Letters on the right indicate brain areas (OL: Optic Lobes, MB: Mushroom Bodies, SIP: Superior Intermediate Protocerebrum, PB: Protocerebral Bridge, FB: Fan-shaped Body, LAL: Lateral Accessory Lobe).

**Extended data figure. 2.**
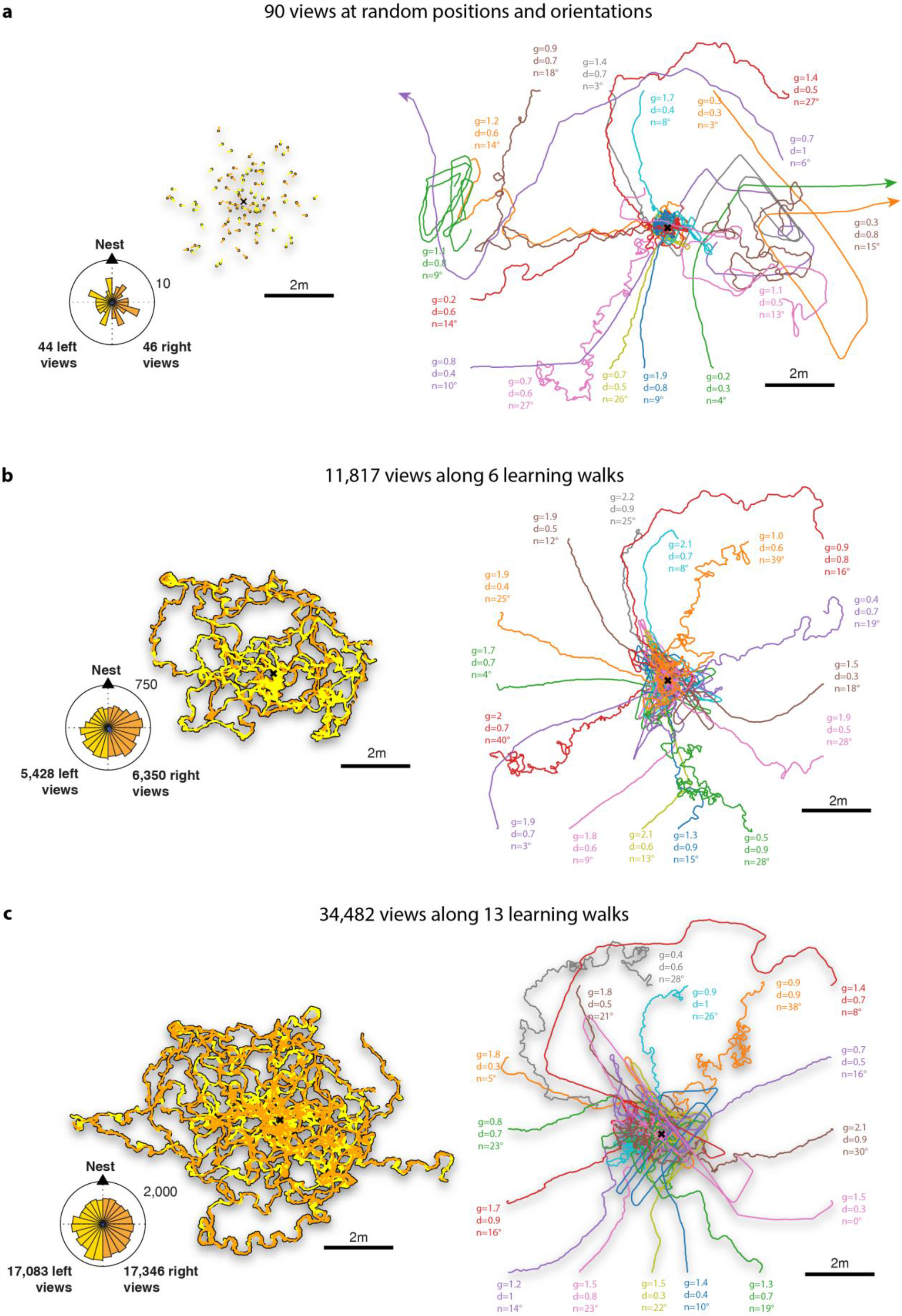
Homing is robust to various training regimes. Paths displayed by the agent when released around the nest with randomly chosen parameter values (g=: motor gain; d=: FBN decay; n=: motor noise, see the ‘parameter description’ section for detailed explanation) (right column) after learning views in different configurations (left column). Orange and yellow indicate how views are categorised as facing right or left from the goal, and thus being respectively learnt in the left or right MB lobes compartments (see Extended data figure 2 for details of the model implementation). Circular histograms show the number of left and right views experienced for learning. **a**. 90 views taken at random positions around the nest (up to 3m away from the nest) and facing in random directions, are enough for the agent to subsequently home and display a search at the nest. The failure of some agents suggests that the catchment area is nonetheless restricted here. **b**,**c**. Using a large amount of views sampled continuously (at 100fps) from multiple real ants learning walks enable the agent to home robustly, and demonstrates that memory load is not a problem. **a**,**b**,**c**. All agents were equipped with 20,000 Kenyon Cells per hemispheres, and embedded in the reconstructed natural world of Canberra (see figure 5), albeit the nest location within the world varied. Note that the scale of movements relative to the world, which can be chosen arbitrarily, is here higher than in figure 5, indicating that the model is effective across various amounts of visual change in relation to movements. This also suggests that the amount of visual change experienced does not need to be precisely controlled by the agent when effecting a learning walk.

**Extended data figure 3.**
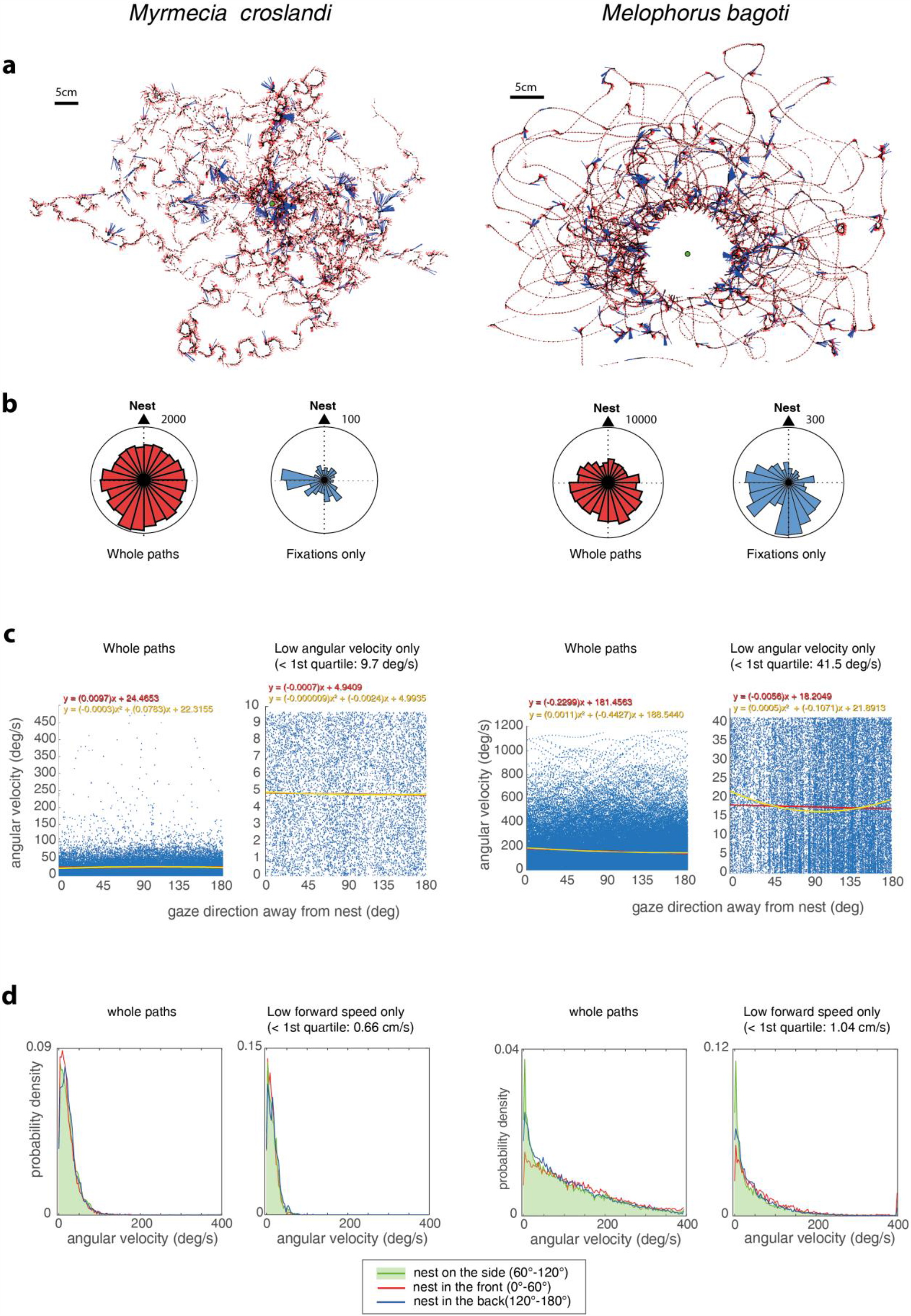
Ants expose their gaze in all direction during learning walks. **a**. Examples of learning walks recorded in *Myrmecia crosslandi* (courtesy of Jochen Zeil, see also ^22^ and *Melophorus bagoti* (from a previous data set used in ^15^) around their nest (green circle). The absence of recording in the centre for M. bagoti results from an experimental funnel around the nest that the ants had to climb before reaching ground level. Dots indicate position of the head (for *M. crosslandi*) or body centroid (for *M. bagoti*) and vectors indicate gaze direction across the recorded frames. Blue vectors mark the instant of ‘fixations’ (when both angular velocity and forward speed are simultaneously < 1^st^ decile of their respective distribution, for each individual). **b**. Circular histogram of the number of views experienced according to their orientation relative to the nest (scale indicated for the circles’ rim), during the whole learning walks or during fixations only (see **a**). Ants show no tendency to bias their gaze towards the nest direction. **c**. Instantaneous angular velocity of the head according to the direction faced relative to the nest, for the whole learning walks or only moments of low angular velocities. In abscises, 0° indicates facing towards the nest, 180° indicates facing in the anti-nest direction, left and right bias are pooled together (by using absolute values). Linear (red) and quadratic fit (yellow) are shown. The flatness of the fits indicates that ants show no tendency to regulate their angular speed according to the direction faced relative to the nest. If anything, in *M. bagoti* ants, low angular speed tends to happen slightly more often when the nest lies on the sides rather than in front or behind. **d**. Relative probability distributions of angular velocities according to whether the nest stands rather in front (0° to 60°, red), in the back (120° to 180°, blue) or on the sides (between 60° and 120°, green line + area). Left and right sides are pulled together using absolute values, so that each of the three categories covers 120° (a third) of the directional space. The similarity of the distributions indicates, here also, no strong tendency to regulate angular speed according to the direction faced relative to the nest; apart from a tendency in *M. bagoti* to display low angular velocities slightly more often when the nest is on the sides.

